# Organic nanoelectronics inside us: charge transport and localization in RNA could orchestrate ribosome operation

**DOI:** 10.1101/2020.08.27.270629

**Authors:** Andrey Sosorev, Oleg Kharlanov

## Abstract

Translation—protein synthesis at the ribonucleic acid (RNA) based molecular machine, ribosome, — proceeds in a similar manner in all life forms. However, despite several decades of research, the physics underlying this process remains enigmatic. Specifically, during translation, a ribosome undergoes large-scale conformational changes of its distant parts, and these motions are precisely synchronized by an unknown mechanism. In this study, we suggest that such a mechanism could be related to charge (electron hole) transport along and between the RNA molecules, localization of these charges at certain sites and successive relaxation of molecular geometry. Thus, we suppose that RNA-based molecular machines, e.g., ribosome, are electronically controlled, having “wires”, “actuators”, “battery”, and other “circuitry”. Taking transfer RNA as an example, we justify the reasonability of our suggestion using ab initio and atomistic simulations. We anticipate that our findings will qualitatively advance the understanding of the key biological processes and could inspire novel approaches in medicine.

## Introduction

For many years after formulation of the central dogma of molecular biology, deoxyribonucleic acid (DNA) has been in the focus of the nucleic acids research, while its “sister molecule” ribonucleic acid (RNA) has been paid much less attention. However, it is now evident that RNAs are not just a bridge between the genetic code stored in DNA and the proteins but instead constitute a key class of molecules that govern or participate in various biochemical processes essential for life of an organism [1–3]. In recent years, non-coding RNAs have attracted particular attention due to importance and variety of their functions. The most studied members of this family—transfer RNAs (tRNAs) and ribosomal RNAs (rRNAs)—enable protein synthesis, or translation, which is a *conditio sine qua non* for an organism to be alive. The state-of-the-art understanding of the process of translation acquired during half a century of research is summarized in several reviews [4–8], however, there is growing evidence that this process is much more complex than it has been assumed for a long time [5].

In all domains of life, translation occurs at the ribosome, which is a complex macromolecular machine built of several RNAs and multiple proteins. This “nanofactory” translates the message encoded in the messenger RNA (mRNA) and synthesizes proteins by linking individual amino acids carried by the cognate tRNAs [5]. Every act of translation includes a number of motions executed by the ribosome and other molecules interacting with it that take place at different ribosome sites. Namely, a ternary complex of aminoacylated tRNA with guanosine triphosphate (GTP) and the so-called translation factor EF-Tu docks into the A-site. If the anticodon of the tRNA matches the codon of the mRNA, the tRNA accommodates, GTP hydrolyzes, EF-Tu is released, and tRNA moves to the P-site, where the reaction of amino acid addition to the growing polypeptide chain takes place. The tRNA then moves to the E-site and finally exits the ribosome from there. Simultaneously with tRNA motion from A to P and finally the E-site, the mRNA is moved by one codon.

Recently, it has been concluded that tRNA motion through the ribosome (translocation) is precisely synchronized with the motion of mRNA [9, 6, 7], large-scale conformational changes of tRNA [10] and ribosome [11–14, 8], and with chemical reactions, namely, GTP hydrolysis in the translation factors [10, 7] and amino acid addition to the growing polypeptide [15]. Importantly, some of these events occur at distant sites of the ribosome separated by several nanometers [16]. For instance, in response to the codon-anticodon match (tRNA “recognition” by the decoding center), the tRNA changes its shape, namely, the anticodon stem gets kinked [17], which results in a correct positioning of the acceptor stem (“accommodation”). The position of this kink—the middle of the anticodon stem [10]—is rather far from the decoding center. Codon-anticodon match also forces the change of the ribosome conformation — the so-called domain closure of its small subunit (SSU) [17, 12]. Further conformational changes alter the global structure of the ribosome, including a swivel of the small subunit head [18, 7], relative rotation of the two ribosomal subunits [19, 7], and movement of the so-called L1 stalk head (by more than 4 nm), which accompanies the tRNA leaving the ribosome [7]. Importantly, these global conformational changes are synchronized [9], although they occur at distant sites of the ribosome [16]: e.g., the distance between the decoding center and the peptidyl-transferase center exceeds 7 nm and that between the L1 stalk and these centers exceeds 9 nm.

Synchronized action of distant parts of the ribosome and tRNAs implies the existence of some mechanism “orchestrating” them [20–22]. Understanding of this mechanism is very important for molecular biology, since it would uncover the physics underlying the key life process; however, this mechanism remains unknown despite several decades of intensive studies of ribosome operation. Several hypotheses on the nature of this mechanism synchronizing the ribosome operation have been suggested, most of them being of mechanistic character [20–22]. Note that such a mechanistic type of allostery (i.e. changes in one site induced by the changes in another, probably distant, site) is frequently observed in proteins [3]. In contrast, one hypothesis attributed the allostery to charge transport along the protein “wires” within the ribosome; moreover, it was suggested that the protein scaffold could operate like a neural network [21]. However, electronic charge transport along proteins is not expected to be efficient since it is hindered by the large distance separating the sites (aromatic residues), between which the charge hops. Nevertheless, there is another route for charge transport in the ribosome—along the RNA helices.

Charge transport in DNA and RNA was discovered more than 50 years ago [23]. Electronic (as opposed to ionic) conductivity of these molecules is enabled by π-stacking of the nucleobases, which can provide significant overlapping of the π-conjugated electron systems of the nucleotides. In this regard, transport in nucleic acids represents the case of an organic (semi)conducting polymer, with polaronic effects and disorder strongly affecting the features of charge migration and localization [24, 25]. At the same time, while several theoretical studies addressed charge transport in DNA, focusing on its applications both in biology and organic electronics [24, 26–28], we are aware of only one such study for RNA [29]. Meanwhile, RNA molecules, especially non-coding RNAs, generally have more complex tertiary structure than DNA [3], allowing for more diverse charge transport topologies. Similarly, while several experimental studies of transport in DNA and its application in active layers of organic electronic devices were performed in the last decade [30–32], for RNA, such studies are rare [33]. Very recently, it has been shown that charge transport along the DNA molecule plays an important role in regulation of the DNA replication [30] and repair [34], and disruption of DNA conductivity may cause serious diseases [35]. Meanwhile, biological role of charge transport along the RNA has not been addressed yet.

In the present study, we propose a general mechanism that could synchronize the motion of distant parts of the ribosome, tRNAs, and mRNA. Specifically, we suggest that this synchronization could be enabled by charge (electron hole) transport along and between the RNA molecules, which results in charge localization at certain sites (areas) of tRNA and, in principle, subsequent conformational changes of the latter. To show this, we model hole transport along a relatively small RNA—phenylalanine tRNA from E. coli (tRNA^Phe^). It turns out that a hole in this molecule can migrate over several nanometers due to efficient overlapping of the π-conjugated systems of the nucleobases. We reveal four sites of charge localization, which are located in the functionally important areas of tRNA. Charge localization is shown to deform both the geometry of these sites and the surrounding nucleotides in the tRNA molecule, which, as we expect, could be related to conformational changes of tRNA during accommodation observed previously. Thus, we suppose that, in a sense, RNA-based machines are electronically controlled like organic electronic devices, with their “conducting wires” transporting charge to “capacitors” that, in turn, power up “actuators”. If our hypothesis is right, the results obtained could contribute to understanding of the basic principles of life on a molecular scale.

## Results

### Site energies, reorganization energies, and transfer integrals

We treat charge transport in tRNA^Phe^ using the hopping model, according to which charge carriers (holes) incoherently hop between the sites—nucleobases (ribose moieties and phosphate chains are not involved in the π-conjugated system and hence are not occupied by the charge carriers). Charge transfer rates are evaluated by the widely used Marcus formula [36], see details in the Methods section. In the Marcus model, the main parameters determining charge migration are (i) the site energies, which are commonly expressed with the frontier orbitals (highest occupied molecular orbital, HOMO, for hole transport; lowest unoccupied molecular orbital, LUMO, for electron transport), (ii) the charge transfer integrals between these sites, *J*, describing their electronic coupling, and (iii) the reorganization energies for sites charging and discharging, *λ*_ch_ and *λ*_dis_, respectively. The HOMO and LUMO energies for the four RNA nucleobases—guanine, adenine, cytosine, and uracil—are depicted in Fig. 1a. As follows from this figure, HOMO energies of the nucleobases are in the range from –7.5 to –5.5 eV, while LUMO energies range from –1 to 0 eV. The high LUMO levels prevent trap-free transport of electrons along the RNA, while moderate HOMO levels provide an opportunity for efficient hole injection and transport in these molecules [37], which was indeed observed in experiments with the “sister” macromolecule, DNA [38]. Therefore, henceforth, we will focus on hole transport (migration) along the RNA. In line with previous findings [39–41], the highest HOMO energy is for guanine; hence, if the holes can migrate along the tRNA, they are expected to localize at guanine bases.

**Fig. 1.**
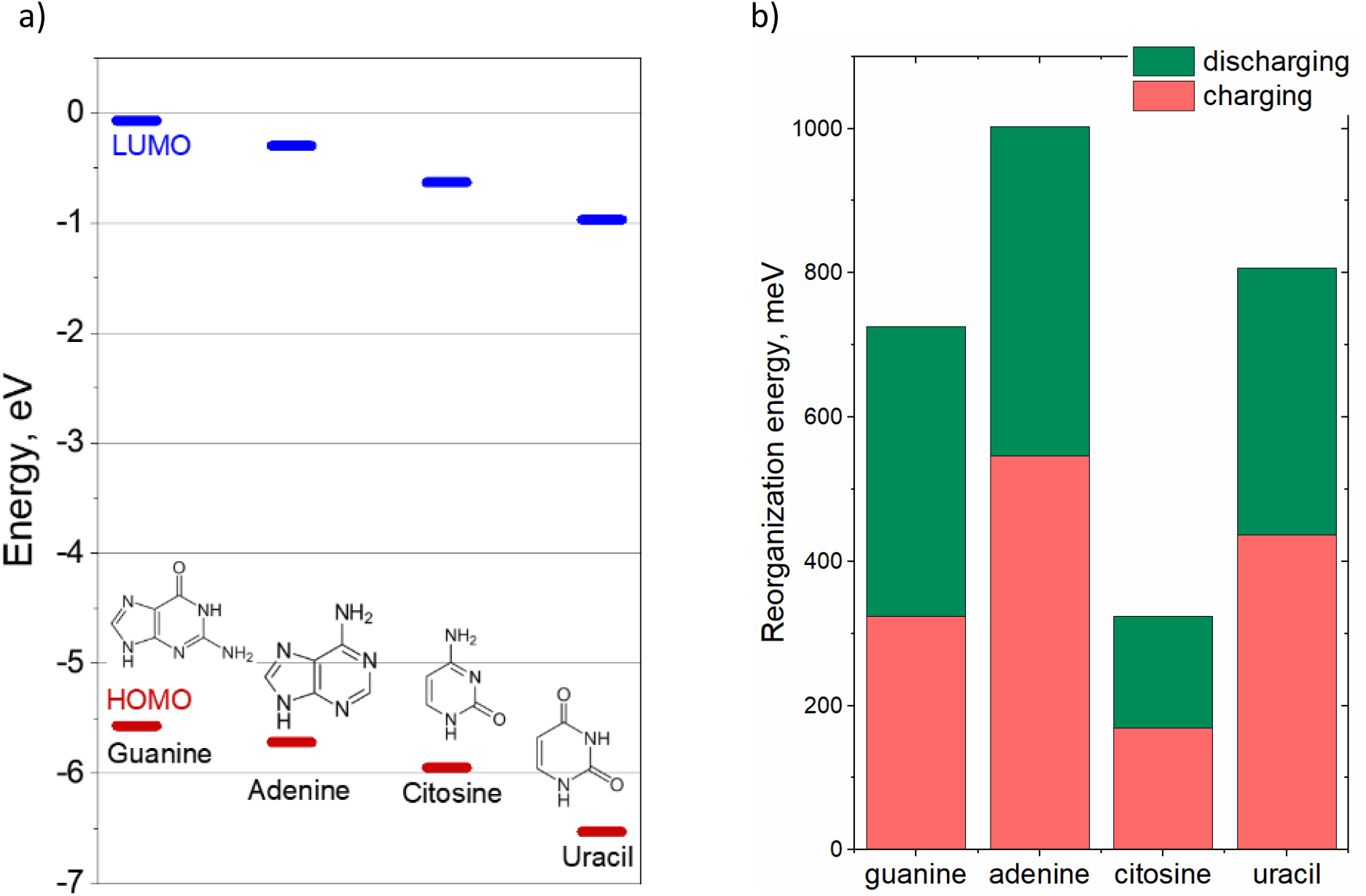
Frontier orbital energies (a) and hole transfer reorganization energies (b) for various nucleobases.

Fig. 1b presents the charging and discharging contributions (*λ*_ch_ and *λ*_dis_, respectively; see Methods section for details) to the total reorganization energy *λ* for each of the four nucleobases. All the *λ* values are large and exceed 300 meV (cf. *λ ∼* 100 meV for organic semiconductors with efficient charge transport, e.g., rubrene and DNTT [42, 43]); the *λ*_ch_ values are comparable to those of *λ*_dis_. Importantly, the *λ* value for guanine—the most probable site for hole localization—is more than 20 times larger than the energy of thermal fluctuations (25 meV at room temperature). This should facilitate the expected hole localization at guanine residues during/after migration: when a hole reaches a guanine residue, the latter undergoes geometry relaxation so that the hole becomes trapped in a deep potential well. Moreover, a high reorganization energy implies a significant geometry relaxation upon charge transfer, as will be shown below.

Fig. 2a illustrates the pattern of transfer integrals *J* for hole transport between various nucleobases of the studied tRNA^Phe^. The *J* values are also listed in Supplementary Information (SI), Table S1. For a number of neighboring nucleobase pairs, the calculated transfer integrals exceed 100 meV. These values are quite large, which is beneficial for charge transport: in many organic semiconductors with high charge mobility, all *J* values are below 100 meV [42, 43]. Inset of Fig. 2a explains the large transfer integrals observed in tRNA: due to the short distance between the nucleobase planes, their HOMO orbitals (responsible for hole transport) overlap significantly, yielding a considerable *J* [44]. Accordingly, Fig. 2b illustrates that there are considerable charge transfer rates *k* for the majority of neighboring bases. In fact, there are two large areas of decent charge transport in the studied tRNA: the acceptor stem with elbow, and the anticodon stem loop with the variable loop. Charge transfer rate between the two areas is negligible. Our calculated rates are comparable to those in DNA reported in Ref. [32].

**Fig. 2.**
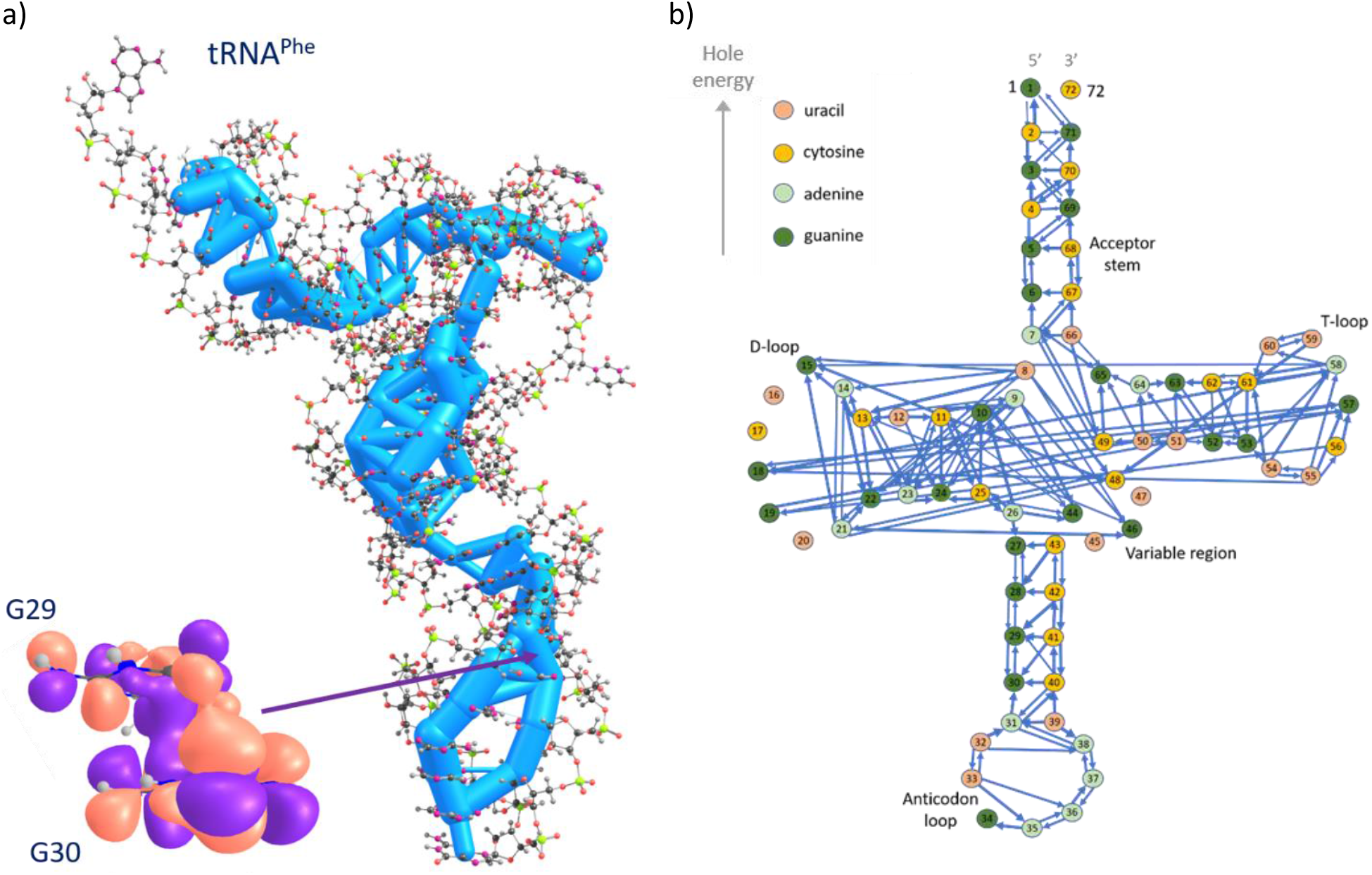
Hole transfer in tRNA^Phe^: (a) transfer integrals *J* and (b) hopping rates *k*. Thicknesses of the cylinders in panel (a) and of the arrows in panel (b) represent the magnitudes of *J* and *k*, respectively, in the logarithmic scale. Inset in panel (a) depicts HOMO delocalization in the guanine dimer (nucleotides G29 and G30; *J*_29,30_ = 69 meV), illustrating strong electronic coupling between the neighboring nucleotides. In panel (b), only rates exceeding 10^−3^ ns^−1^ are shown.

### “The wires and capacitors” : Charge migration and localization

To address the pathways of hole migration across the charge transporting network of tRNAs, we performed a Kinetic Monte Carlo simulation: a hole was sequentially placed at various initial sites, then its motion was monitored, and probabilities to find the hole at various final sites were calculated. Fig. 3a presents the transition probabilities to find the hole, which started migration from the given initial site, at the given final site. This figure clearly reveals a localization pattern of the charge carrier. Namely, vertical cyan-green lines indicate the sites of charge localization: the corresponding final sites collect holes from many initial sites; the longer the lines, the larger the “basins of attraction”. Final populations, i.e. the probabilities to find an initially randomly placed hole at a given site, are shown in Fig. 3b. As follows from Figs. 3a,b, holes most often localize at the four nucleotides G27, G10, G34, and G63. As expected, all of them are guanines. Localization becomes more pronounced with time; the time dependence of the localization pattern is presented in Supplementary Fig. S2. Moreover, addition of a static Gaussian energy disorder with an amplitude of 25 meV does not smear out or even considerably alter the localization picture in Fig. 3b (see Supplementary Fig. S3a), even though the spread of HOMO energies within tRNA^Phe^ is also 25–30 meV for each of the four nucleobases. Static disorder of transfer integrals with a scale *σ*_*J*_ ∼ |*J*| previously demonstrated for DNA [24] does not produce any effect either, at least at the level of Marcus kinetics (see Supplementary Fig. S3b). Last but not least, restricting the initial hole position to the anticodon stem-loop (G27–C43 nucleotides) drastically changes the probabilities in Fig. 3b: holes from the whole stem end up at G27 with a 43% probability (see Fig. 3c and, regarding robustness against disorder, Supplementary Fig. S3a,b).

**Fig. 3.**
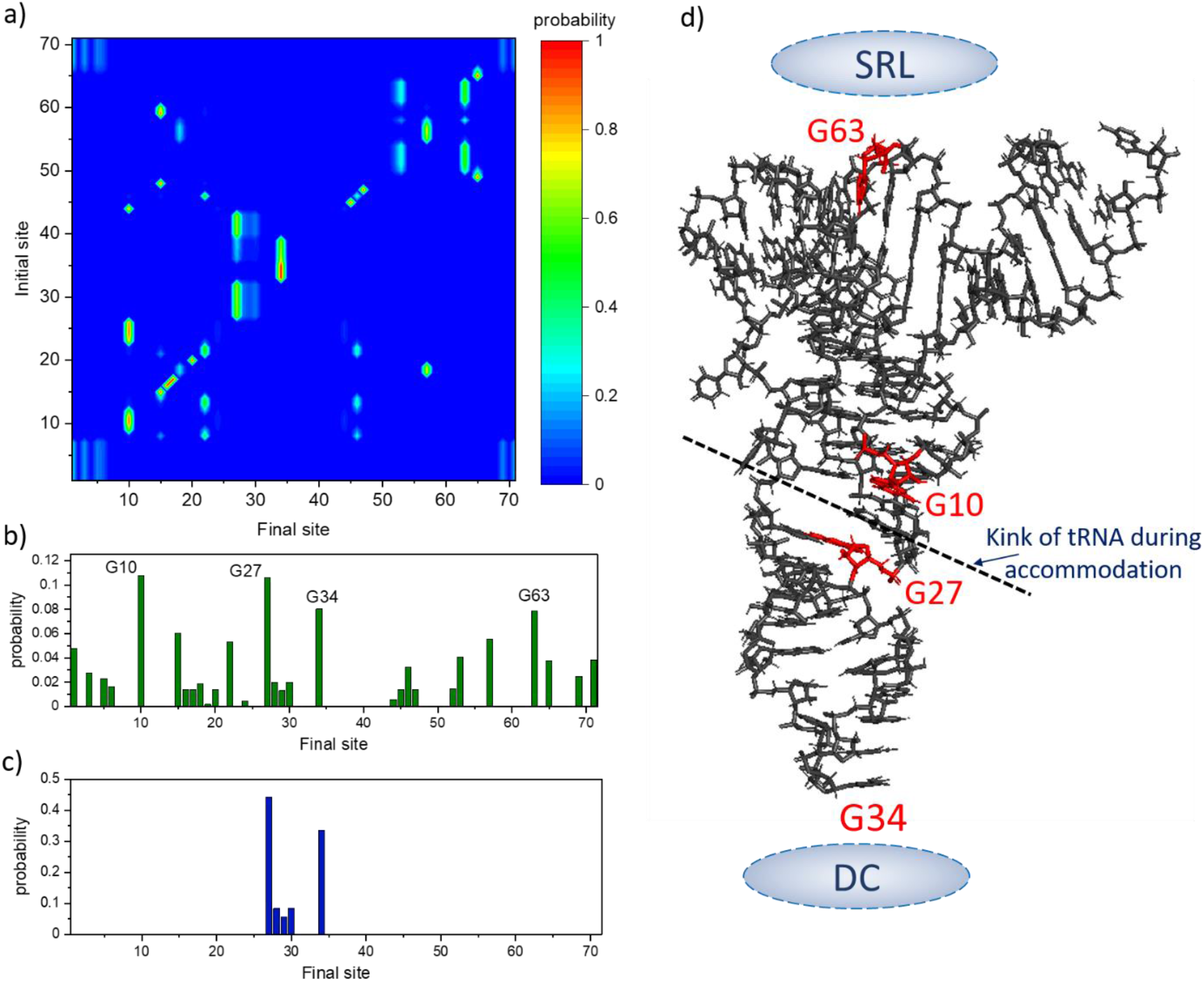
Localization of holes in tRNA. (a) Transition probabilities of finding a hole, initially placed at a given site, at different final sites at the moment *t* = 500 ns. (b) Average populations of different final sites, i.e. percentages of the initial sites, starting from which the hole localizes at a given final site, obtained via summing the transition probabilities over the initial sites. (c) Average final populations for a hole initially placed at the anticodon stem-loop G27–C43 nucleotides. (d) Four sites with the most probable hole localization from panel (b). In panel (d), the position of the tRNA kink during accommodation (dashed line), the sarcin-ricin loop (SRL) of rRNA (top gray ellipse) and decoding center (DC, bottom gray ellipse) are also sketched.

Notably, the abovementioned localization sites are situated in the functionally important areas of tRNA, as sketched in Fig. 3d. Specifically, G27 and G10 are adjacent to the origin of the anticodon stem-loop kink that occurs around position 26 during tRNA accommodation [45]. G63 is in close contact with the sarcin-ricin loop (SRL) of the rRNA. In turn, SRL approaches the GTP domain of EF-Tu during tRNA accommodation, and we suggest that a hole localized at G63 or SRL could, in principle, trigger rearrangement of the EF-Tu structure, finally resulting in GTP hydrolysis at the appropriate moment. G34 is located in the anticodon, near the decoding center (DC).

### “The actuators” : charge-localization-induced conformational changes

Since charge carriers can migrate along tRNA (and rRNA, see below) and localize in their functionally important areas, this remarkable feature is likely to play a certain role in ribosome operation. As mentioned above, we suggest that this role should be related to regulation of the translation process. A possible mechanism underlying this regulation is the changes of the molecular conformation resulting from charge localization. Indeed, charge localization at certain sites of tRNA (see Fig. 3d) inevitably results in deformation of the corresponding nucleotides and alteration of their arrangement. Fig. 4 illustrates these changes for a G nucleotide and a G-C pair found from DFT simulations and demonstrates the effect of a hole localized at G27 on the geometry of the anticodon stem-loop. Let us discuss these induced deformations in more detail.

**Fig. 4.**
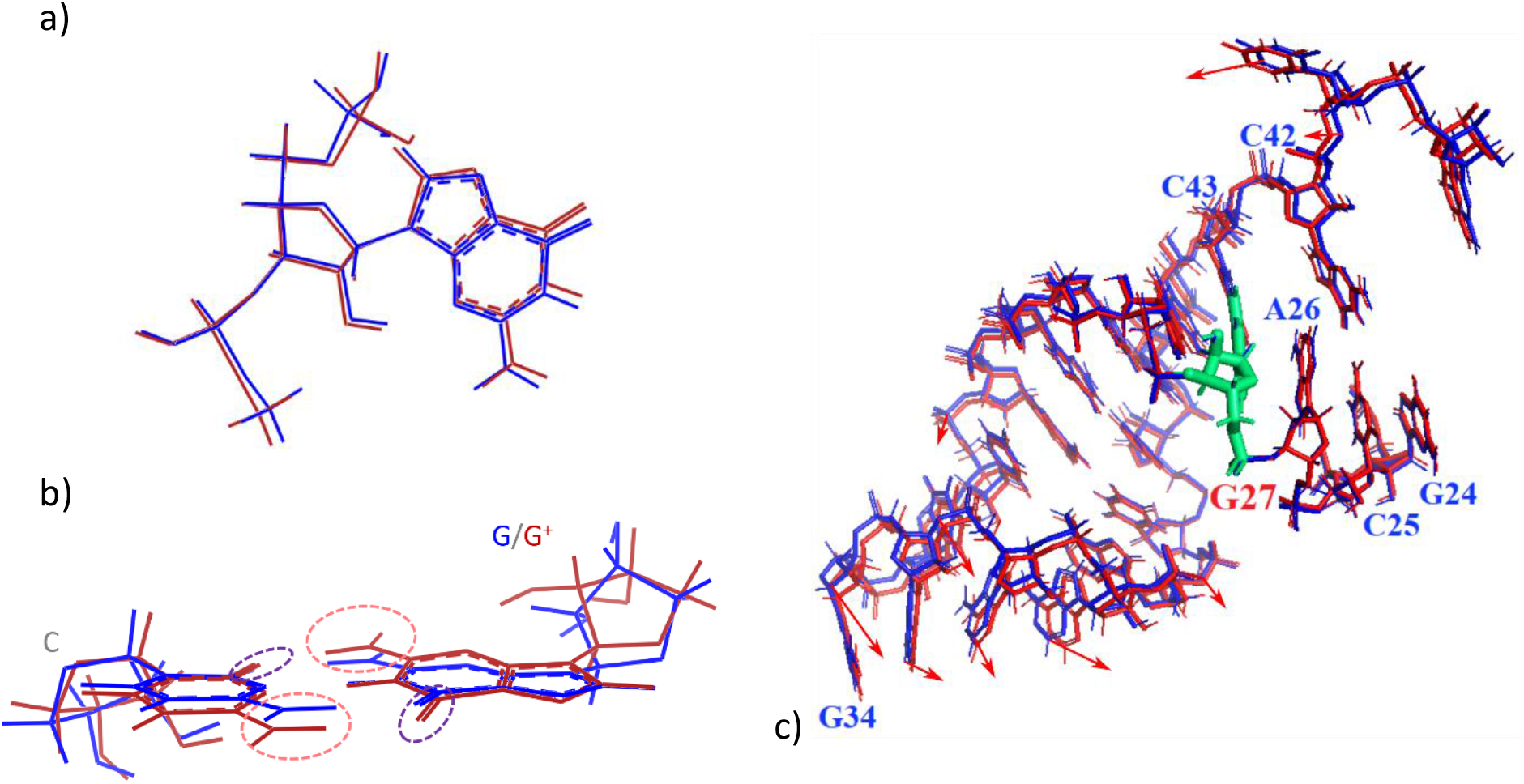
Conformational changes of tRNA molecule and its building blocks induced by the presence of a localized positive charge (the neutral configuration in blue versus the charged configuration in red). (a) Geometry change of a guanine residue (DFT). (b) Deformation of a guanine-cytosine pair (DFT). Carbonyl group is highlighted with a dashed violet ellipse, and amino group is highlighted with a dashed pink ellipse. (c) Geometry changes taking place in the anticodon stem-loop of a tRNA^Phe^ molecule due to a hole localized at the (green) G27 residue (molecular mechanics, geometry relaxation in explicit solvent). Red arrows denote directions and relative magnitudes of residue displacements in a hole-hosting tRNA, with respect to a neutral one.

As follows from Fig. 4a, the positively charged guanine nucleobase approaches the ribose moiety and phosphate chain, likely because of the presence of negatively charged areas at the latter one (see Supplementary Fig. S4b). Moreover, Fig. 4b shows that the relative orientation of the charged guanine nucleotide G27 with respect to its partner in pair (cytosine C43) changes as well: the positively charged amino group (which increases its positive charge in the cation state, see Supplementary Fig. S5) approaches the negatively charged carbonyl group of the partner, while its own carbonyl group (which decreases its negative charge in the cation state) moves away from the positively charged amino group of the partner. Thus, the sites of charge localization can serve as nano-actuators: upon gaining a hole, they change their conformation. Importantly, the changes in the geometry induced by charge localization are stable and can hardly be disrupted by thermal motion, since the reorganization energy (∼700 meV for guanine) substantially exceeds the energy of thermal fluctuations (∼25 meV).

Now, although local atomic displacements shown in Fig. 4a,b do not exceed 1 Å, they are expected to be enhanced by the adjacent moieties, as it is observed in proteins, e.g., EF-Tu upon GTP hydrolysis [3]. Indeed, Fig. 4c demonstrates that the effect of a hole hosted by the G27 residue of tRNA propagates to the whole anticodon stem-loop (i.e. down to the G34 residue), leading to its slight bending towards G27. It should be noted here that this result was found using molecular mechanics, while a careful analysis of the global tRNA conformation requires a time-consuming molecular dynamics (MD) simulation in explicit solvent, which we leave beyond this study. However, localized charges can induce large-scale conformational changes of nucleic acid molecules at level of MD (e.g., for DNA, see Ref. [46]). Noteworthily also, guanine tracts (which are the most favorable places for hole localization and thus are possibly related to conformational changes) are present in anticodon stems of several other tRNAs – e.g., mouse selenocysteine tRNA and arginine, cysteine, glutamine, glycine and other tRNAs from E. coli. Finally, it is worth noting that the conformational changes are the usual mechanism underlying the allostery in various proteins [3].

### “The battery and power cord” : the source of charge carriers and their pathway to tRNA

The presented scenario of translation regulation via hole transport, localization, and possible subsequent geometry changes lacks an explanation where the holes come from and where they leave. We reckon that the role of the “battery” that provides holes and then probably takes them away could be played by the Fe_4_S_4_ clusters, as was earlier proposed for DNA replication and repair [30, 47]. Indeed, these redox-active clusters that can readily provide or gain electrons/holes without considerable reorganization [47] have been recently discovered in the small subunit of the ribosome [48]. However, the very role of the Fe_4_S_4_ cluster in the ribosome has not been identified [48]. In light of our scenario, it seems possible that the cluster could provide holes migrating towards certain functional sites and inducing conformational changes that are required for the synchronized operation of the parts of the molecular machine. Indeed, the charge transport should not be limited only to tRNA but can also occur in rRNA since the latter forms long helices, in which nucleotides are well stacked, implying efficient charge transport [50]. Fig. 5 clearly shows such a charge transport pathway from the Fe_4_S_4_ cluster to the tRNA anticodon through rRNA. In the latter one, efficient charge transport is expected due to considerable transport integrals (see Supplementary Fig. S6), like in the tRNA studied herein.

**Fig. 5.**
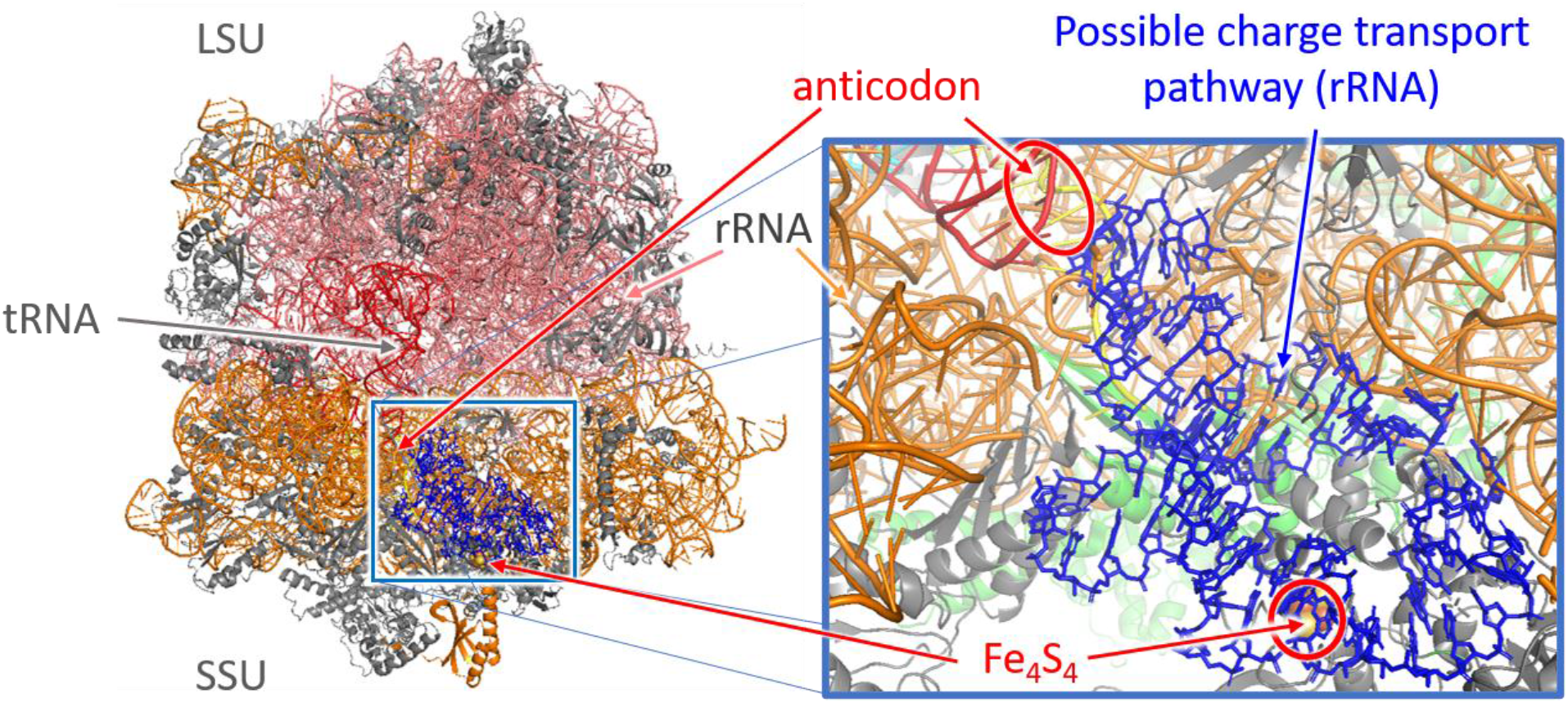
A possible charge transport path between Fe_4_S_4_ cluster of the small subunit (SSU) of the ribosome and the tRNA anticodon. LSU stands for the large subunit of the ribosome. The geometry corresponds to PDB entry 6QNQ [48, 49].

## Discussion

This study addressed the biological role of charge transport along the RNA molecules for the first time. Being primarily aimed at presentation and justifying the relevance of our scenario of translation “orchestration”, it does not pretend on a quantitatively accurate description of the charge transport process; this is why the methods we used here were rather simple. Specifically, we followed a simple multiscale modelling scheme combining DFT calculations with molecular mechanics, whereas a more rigorous analysis of the discussed phenomena would require a kind of time-consuming ab-initio molecular dynamics. Also, we intentionally used a rather simple charge transport model based on the Marcus theory, although charge transport in nucleic acids can be more complex and still remains a subject of debates [40]. Possible charge delocalization between the sites that could decrease the effective reorganization energies [51–53, 43] was neglected. Moreover, the very site energies (ionization potentials of nucleotides) can be dependent on charge delocalization [54] and hence on *J*. Ionic conductivity, which can be coupled to the electronic charge transport discussed herein [55], is also accounted for simply at the level of counterions added to tRNA. Finally, electronic transport discussed herein could trigger proton transport, which was recently suggested to occur in the peptidyl-transferase center of the ribosome [56]. Nevertheless, we do believe that accounting for the abovementioned factors will not question the qualitative side of our conclusions but will rather refine the model suggested herein.

Although in this study we focused on the role of charge transport in tRNA accommodation, the “charge transport–localization–conformational changes” scenario could, in principle, synchronize other stages of translation elongation, as well as that of translation initiation and termination. For instance, we suppose that Fe_4_S_4_ clusters can serve not only for charge donation/acquisition, but also as a “ratchet” controlling relative rotation of ribosome subunits during translocation [8, 9]. Indeed, it was shown in Ref. [47] that proteins bearing oxidized Fe_4_S_4_^3+^ are ∼500 times more strongly bound to DNA than those bearing an equilibrium-state Fe_4_S_4_^2+^. By analogy, upon hole acquisition by a Fe_4_S_4_ cluster of uS4 protein located in the SSU [48], the ribosome could become “locked”, while after discharging the Fe_4_S_4_, the ribosome could “open”. Moreover, the possible “ratcheting” role of the hole localization is not necessarily associated only with the Fe_4_S_4_ cluster: the hole could localize in other areas, probably, at the interface between the SSU and LSU (intersubunit bridges), and affect the relative motion of the subunits. The suggested scenario could be also responsible for synchronization of other large-scale conformational changes of the ribosome—SSU head swiveling during tRNA accommodation and L1 stalk motion during the tRNA departure from the E-site—which are correlated with other steps of the translation process [9]. Finally, Fe_4_S_4_ clusters are observed not only in the ribosome, but also in protein co-factor involved in translation initiation [57], translation termination [58], and ribosome recycling [57] in eukaryotes. Since biosynthesis of Fe_4_S_4_ clusters is rather expensive [47], their presence in the described systems should play an important role in the operation of these systems, e.g., they could serve as “batteries” providing charges for electronic control of translation initiation, termination, and ribosome recycling. Moreover, the clusters from this factor and ribosome can communicate via charge transport along RNA, by analogy with the hypothesis suggested in Refs. [30, 47] for DNA-mediated communication.

We anticipate that the proposed “charge transport–localization–conformational changes” mechanism can also synchronize the operation of other RNA-based molecular complexes (machines), e.g., signal recognition particle RNAs, spliceosomes, and other ribozymes, which undergo large-scale conformational changes during their operation. Moreover, our result could clarify the key stages of the operation of viruses, viroids, and virus-associated RNAs. RNAs in these systems are also arranged in stems and helices and can switch between various conformations in dependence of the desired action. For instance, it was discovered that changes in the conformation of one of the two domains in 3’ untranslated region (UTR) of turnip yellow mosaic virus RNA induced conformational changes in the other domain separated by a rather long helix [59]. The authors of Ref. [59] concluded that some conduit existed between them, but the nature of this interconnection was not established. We suggest that the latter could well work according to the “charge transport–localization–conformational changes” scenario.

Extending the analogy with the conventional electronic devices, one is thus able to say that some sites of RNA-protein complexes, e.g., the ribosome, can serve as nanotransistors. Specifically, at these sites, charge transport along the RNA (“source-drain current”) can be modulated by a protein (“gate”) that changes its conformation in response to some external signal. These nanotransistors could perform elementary logical operations, which would explain the “smartness” of these complexes in performing their functions. Noteworthily, understanding of the operation of RNA-based molecular machines can facilitate formulation of new approaches to treatment of various diseases. Specifically, it could help with discovering antiviral drugs (especially against RNA viruses) and anticancer ones (since it is now assumed that several long non-coding RNAs participate in cancer progression [60])—two very challenging issues of the modern medicine. Certainly, such issues need to be thoroughly investigated theoretically and experimentally, in order to get beyond the level of far-reaching hypotheses inspired by a general scenario, and are a subject or our ongoing studies.

To conclude, we have suggested a physical mechanism possibly underlying synchronization of the large-scale conformational changes of RNA molecules occurring during protein synthesis at the ribosome. Taking tRNA as an example, we have shown that RNA molecules can efficiently conduct electron holes, which localize at very specific molecular sites (nucleobases) and are able to affect the geometry of the molecule. Importantly, for the tRNA studied, such sites are situated in its functionally important areas, related to the key processes underlying translation, such as the kink of the tRNA anticodon stem-loop that is necessary for tRNA accommodation. The proposed “charge transport–localization–conformational changes” mechanism can at least partially explain the correlated motion of distant ribosome parts during translation, and perhaps the operation of other RNA-based molecular machines as well. The latter thus resemble macroscopic electronically controlled machines and contain their “batteries”, “wires”, and “actuators”. Our findings are anticipated to improve understanding of the physical mechanisms underlying the key processes taking place in the cells, advance the molecular biology, and open up a way towards new approaches to treatment of diseases.

## Methods

**Initial geometry** of the tRNA studied was extracted from a PDB database (entry 3L0U) [61, 49]. Then it was relaxed using AMBER94 force field [62] within GROMACS package [63, 64]. For this purpose, the RNA structure was placed in a cubic water box with the size equal to the largest distance between the atoms of the considered system plus 2 Å. All density functional theory (DFT) calculations were performed in GAMESS package [65, 66] using CAM-B3LYP (for reorganization energies) or B3LYP (otherwise) density functionals and 6-31g(d) basis set.

**Charge transport** was simulated using the hopping model based on the Marcus formula for charge transfer rate from site *a* to site *b* [36]:

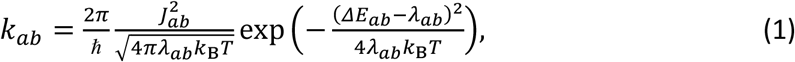

where *ℏ* is the reduced Planck constant, *k*_B_ is the Boltzmann constant, *T* is the absolute temperature, *J*_*ab*_ is the charge transfer integral between sites *a* and *b, λ*_*ab*_ is the reorganization energy for charge transport between these sites (the sum of the relaxation energies for hole loss by site *a, λ*_*a*,dis_, and hole gain by site *b, λ*_*b*,ch_, see below), and *ΔE*_*ab*_ = *E*_*a*_ – *E*_*b*_ is the HOMO energy difference between the initial and the final sites. For the calculation of HOMO energies for various sites (nucleotides), *E*_*a*_, and hole transfer integrals, *J*_*ab*_, the nucleotides were substituted by nucleobases with methyl groups instead of ribose moiety. *E*_*a*_ were calculated for the nucleobase geometries obtained from molecular mechanics geometry optimization of tRNA. Transfer integrals *J*_*ab*_ were calculated using the dimer projection method (DIPRO) [67–69]. Calculations were performed in vacuum; accounting for water using the polarizable continuum model altered *J*_*ab*_ values by less than 20%. For the calculation of reorganization energies, *λ*_*ab*_, the nucleotides were substituted by nucleobases with hydrogens instead of ribose moiety. The *λ*_*ab*_ values were approximated by their internal (intramolecular) parts and calculated according to the standard adiabatic potentials (four-point) scheme [44, 53]. Specifically, *λ*_*ab*_ was calculated as *λ*_*ab*_ = *λ*_*b*,ch_ + *λ*_*a*,dis_ = (*E*_N*b*_ ^*\**^ – *E*_N*b*_) + (*E*_C*a*_ ^*\**^ – *E*_C*a*_), where *E*_N*b*_ and *E*_N*b*_ ^*\**^ are the energies of the neutral state of site *b* in its optimized geometry and in the optimized geometry of the charged state, respectively, while *E*_C*a*_ and *E*_C*a*_ ^*\**^ are the energies of the charged state of site *a* in its optimized geometry and in the geometry of the neutral state, respectively.

**Molecular mechanics** simulations were performed in AmberTools 16 package, with explicit water solvent and sodium counterions added to the tRNA molecule. A standard AMBER RNA force field ff99 + bsc0 + *χ*OL3 [70] was used for tRNA, while the solvent was described by the TIP3P model [71]. Excess positive charge localized on a given nucleotide was introduced into the force field via shifting the atomic charges *q*_*i*_ in accordance with DFT calculations of the nucleobases: namely, AMBER charges of atoms *i* belonging to the G27 nucleobase were shifted by a difference between the Mulliken charges of the corresponding atoms of the guanine nucleobase in its charged and neutral states, 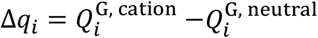 (see Supplementary Fig. S5). For visualization of results, ChemCraft, PyMOL, and Jmol software were used.

## Acknowledgements

The work was supported by Russian Foundation for Basic Research (project No. 19-32-60081). The authors are grateful to D. Paraschuk and R. Efremov for valuable discussions and to I. Chernyshov for initial optimization of tRNA geometry using molecular mechanics.

