## Supplementary Information for "Organic nanoelectronics inside us: charge transport and localization in RNA could orchestrate ribosome operation"

### S1. Hole transfer integrals and transfer rates

**Table S1** Hole transfer integrals between various sites (only those exceeding 1 meV are presented)

| Site pairs *a–b* | Site types | $J_{ab}$, eV | Site pairs *a–b* | Site types | $J_{ab}$, eV | Site pairs *a–b* | Site types | $J_{ab}$, eV | Site pairs *a–b* | Site types | $J_{ab}$, eV |
|---|---|---|---|---|---|---|---|---|---|---|---|
| 1–2 | G–C | 0.156 | 10–24 | G–G | 0.005 | 26–44 | A–G | –0.01 | 50–64 | U–A | 0.015 |
| 1–71 | G–G | 0.007 | 10–25 | G–C | –0.023 | 27–28 | G–G | 0.007 | 50–65 | U–G | –0.027 |
| 2–3 | C–G | –0.012 | 10–26 | G–A | 0.111 | 27–43 | G–C | –0.015 | 51–52 | U–G | –0.002 |
| 3–4 | G–C | 0.136 | 10–44 | G–G | 0.012 | 28–29 | G–G | 0.001 | 51–63 | U–G | 0.008 |
| 3–69 | G–G | 0.001 | 11–12 | C–U | 0.045 | 28–42 | G–C | –0.023 | 51–64 | U–A | –0.004 |
| 3–70 | G–C | –0.021 | 11–23 | C–A | 0.003 | 28–43 | G–C | –0.049 | 52–53 | G–G | –0.079 |
| 3–71 | G–G | 0.086 | 11–24 | C–G | –0.011 | 29–30 | G–G | –0.069 | 52–63 | G–G | 0.052 |
| 4–5 | C–G | –0.004 | 11–25 | C–C | 0.017 | 29–41 | G–C | 0.004 | 53–54 | G–U | –0.191 |
| 4–69 | C–G | 0.002 | 12–13 | U–C | –0.036 | 29–42 | G–C | –0.055 | 53–58 | G–A | –0.003 |
| 4–70 | C–C | 0.059 | 12–23 | U–A | –0.026 | 30–31 | G–A | 0.005 | 53–61 | G–C | –0.024 |
| 5–6 | G–G | –0.071 | 12–24 | U–G | 0.013 | 30–40 | G–C | –0.02 | 53–62 | G–C | –0.018 |
| 5–68 | G–C | –0.023 | 13–14 | C–A | 0.019 | 30–41 | G–C | 0.016 | 54–55 | U–U | 0.105 |
| 5–69 | G–G | 0.091 | 13–22 | C–G | 0.014 | 31–32 | A–U | –0.08 | 54–58 | U–A | 0.006 |
| 6–7 | G–A | –0.033 | 13–23 | C–A | –0.03 | 31–38 | A–A | 0.014 | 54–61 | U–C | –0.007 |
| 6–67 | G–C | –0.017 | 14–15 | A–G | –0.067 | 31–39 | A–U | –0.013 | 55–56 | U–C | –0.003 |
| 6–68 | G–C | –0.058 | 14–21 | A–A | –0.003 | 31–40 | A–C | 0.071 | 55–57 | U–G | 0.064 |
| 7–49 | A–C | –0.044 | 14–22 | A–G | –0.13 | 32–33 | U–U | 0.068 | 55–58 | U–A | –0.002 |
| 7–65 | A–G | –0.002 | 15–21 | G–A | 0.015 | 32–38 | U–A | 0.001 | 56–57 | C–G | –0.039 |
| 7–66 | A–U | –0.017 | 15–48 | G–C | 0.008 | 33–35 | U–A | 0.04 | 58–60 | A–U | –0.001 |
| 7–67 | A–C | 0.06 | 15–59 | G–U | 0.001 | 33–36 | U–A | –0.002 | 58–61 | A–C | 0.139 |
| 8–13 | U–C | 0.069 | 18–55 | G–U | 0.006 | 34–35 | G–A | 0.014 | 59–60 | U–U | –0.102 |
| 8–14 | U–A | –0.005 | 18–57 | G–G | 0.049 | 35–36 | A–A | –0.09 | 60–61 | U–C | 0.003 |
| 8–15 | U–G | 0.003 | 18–58 | G–A | –0.011 | 36–37 | A–A | 0.134 | 61–62 | C–C | –0.001 |
| 8–21 | U–A | 0.001 | 19–56 | G–C | –0.002 | 37–38 | A–A | 0.071 | 62–63 | C–G | –0.006 |
| 8–22 | U–G | –0.003 | 19–57 | G–G | 0.081 | 38–39 | A–U | 0.104 | 63–64 | G–A | 0.04 |
| 8–46 | U–G | 0.002 | 21–22 | A–G | 0.022 | 39–40 | U–C | –0.012 | 64–65 | A–G | –0.001 |
| 8–48 | U–C | 0.031 | 21–46 | A–G | –0.035 | 40–41 | C–C | 0.016 | 66–67 | U–C | 0.017 |
| 9–11 | A–C | 0.004 | 21–48 | A–C | –0.02 | 41–42 | C–C | –0.018 | 67–68 | C–C | –0.049 |
| 9–22 | A–G | 0.043 | 22–23 | G–A | 0.058 | 42–43 | C–C | –0.003 | 68–69 | C – G | –0.03 |
| 9–23 | A–A | –0.019 | 22–46 | G–G | –0.007 | 48–59 | C–U | –0.079 | 69–70 | G – C | 0.039 |
| 9–24 | A–G | –0.004 | 23–24 | A–G | –0.12 | 49–50 | C–U | –0.013 | 70–71 | C – G | –0.028 |
| 9–44 | A–G | 0.055 | 24–25 | G–C | –0.011 | 49–65 | C–G | –0.014 | | | |
| 9–46 | A–G | –0.004 | 25–26 | C–A | 0.029 | 49–66 | C–U | –0.02 | | | |
| 10–11 | G–C | 0.033 | 26–27 | A–G | 0.022 | 50–51 | U–U | –0.095 | | | |

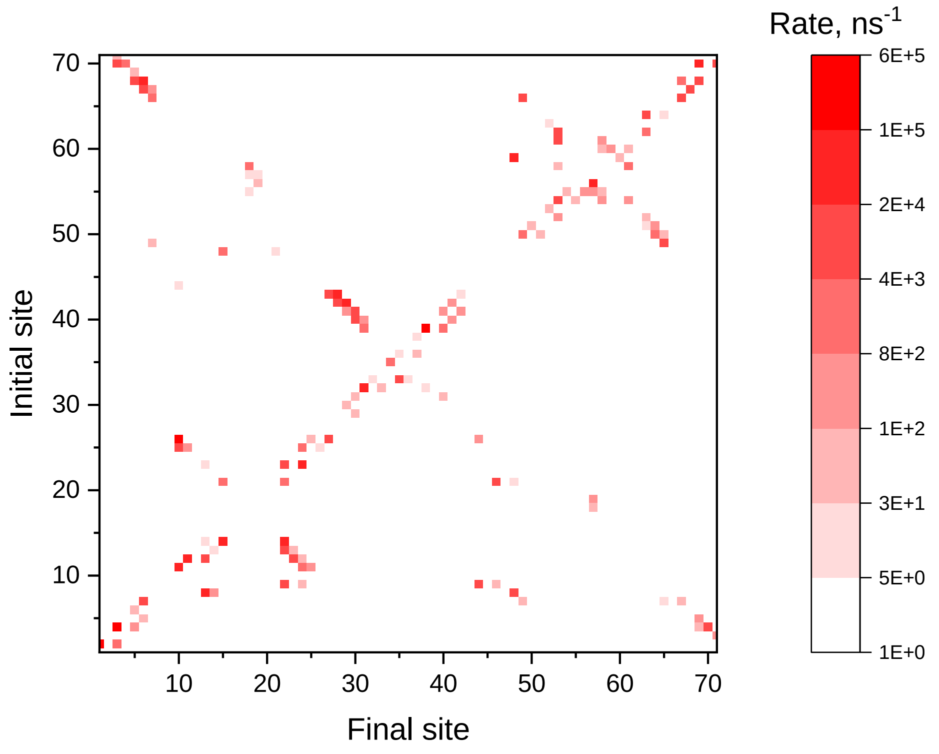


**Fig. S1** Hole transfer rates $k_{ab}$ between various sites of $tRNA^{Phe}$, in $ns^{-1}$

### S2. Dynamics of charge localization

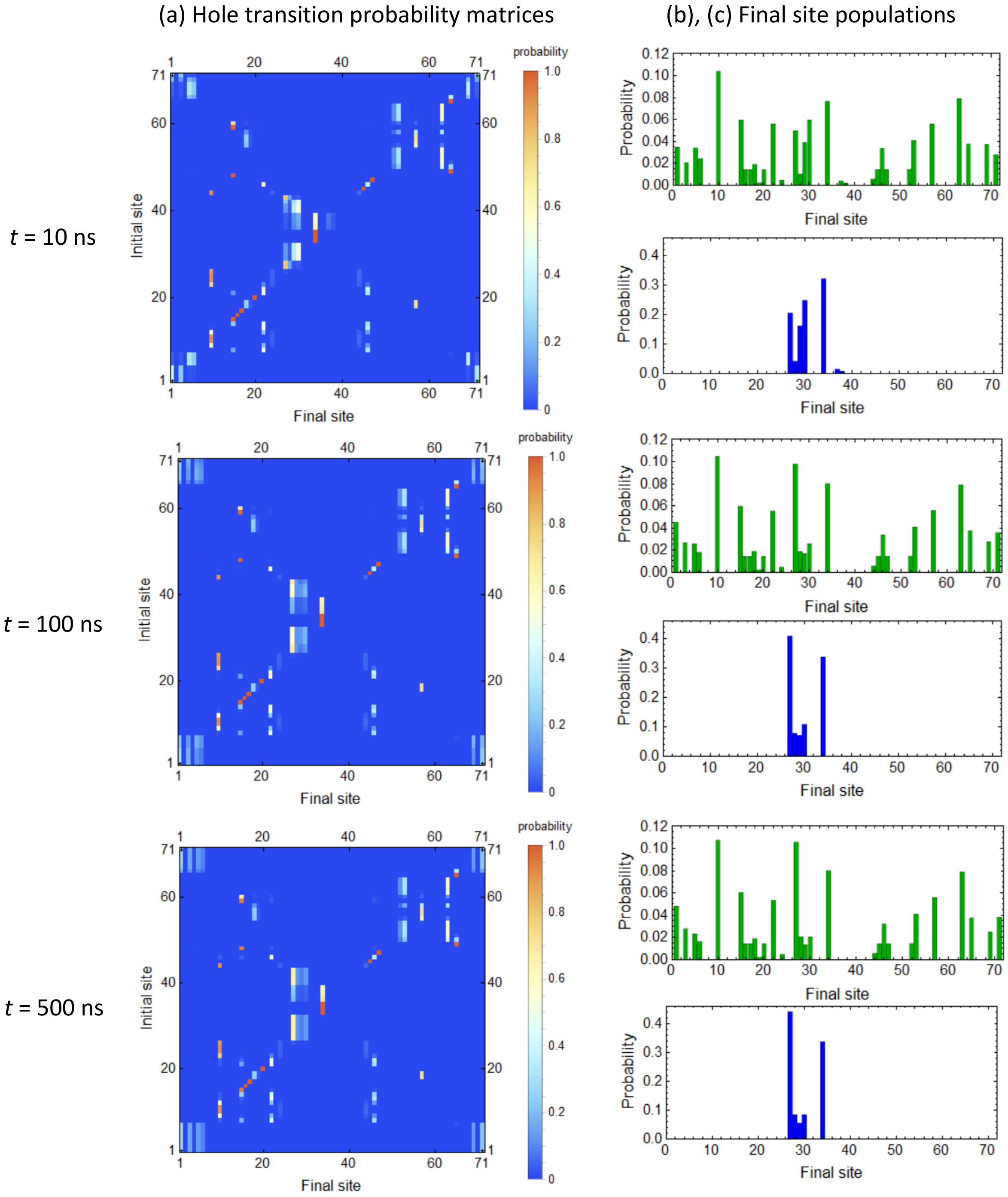


**Fig. S2** Dynamics of hole localization pattern in tRNA$^{Phe}$: (a) transition probability matrices and (b), (c) site (nucleobase) populations at different moments of time. Green and blue bars in panels (b), (c) demonstrate populations for a hole starting from an arbitrary site of the whole tRNA and from an arbitrary site within the anticodon stem-loop, respectively

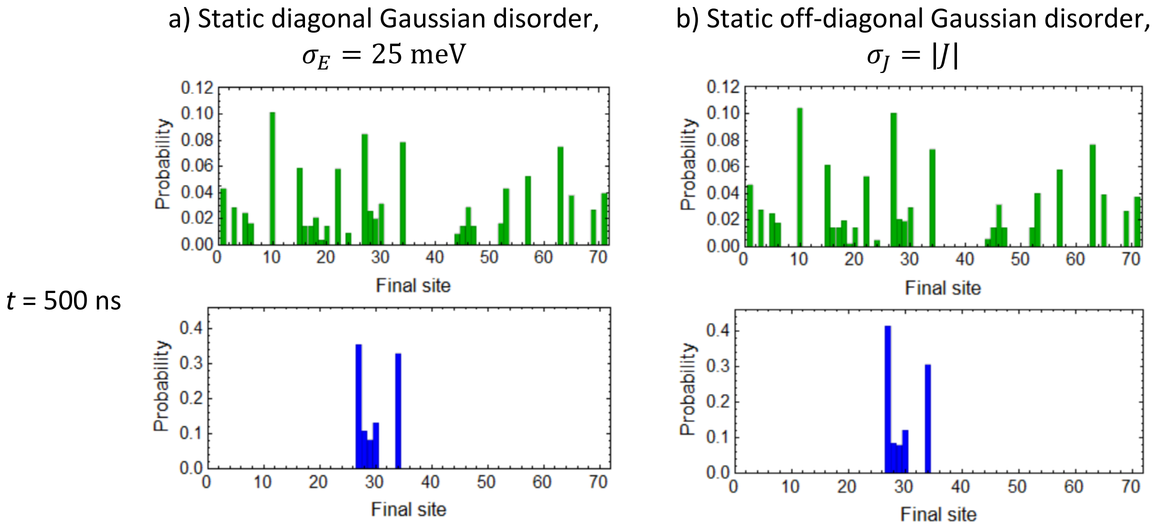


**Fig. S3** Effect of Gaussian static disorder on hole localization in tRNA$^{Phe}$. Green and blue bars demonstrate populations for a hole starting from an arbitrary site of the whole tRNA and from an arbitrary site within the anticodon stem-loop, respectively

### S3. Electrostatic potential of nucleobases

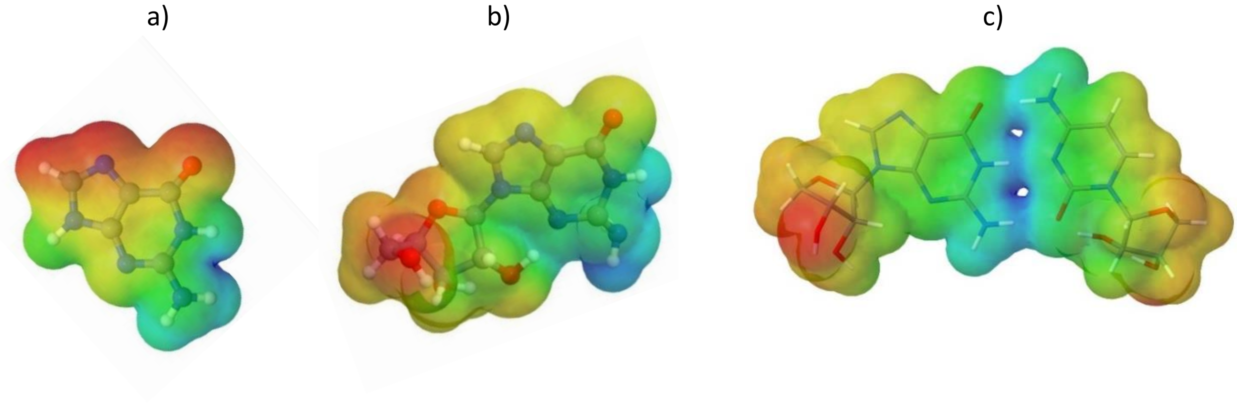


**Fig. S4** Electrostatic potential of guanine (a), guanosine (b) and G-C pair (c). Red color stands for negative electrostatic potential, and blue color stands for positive electrostatic potential

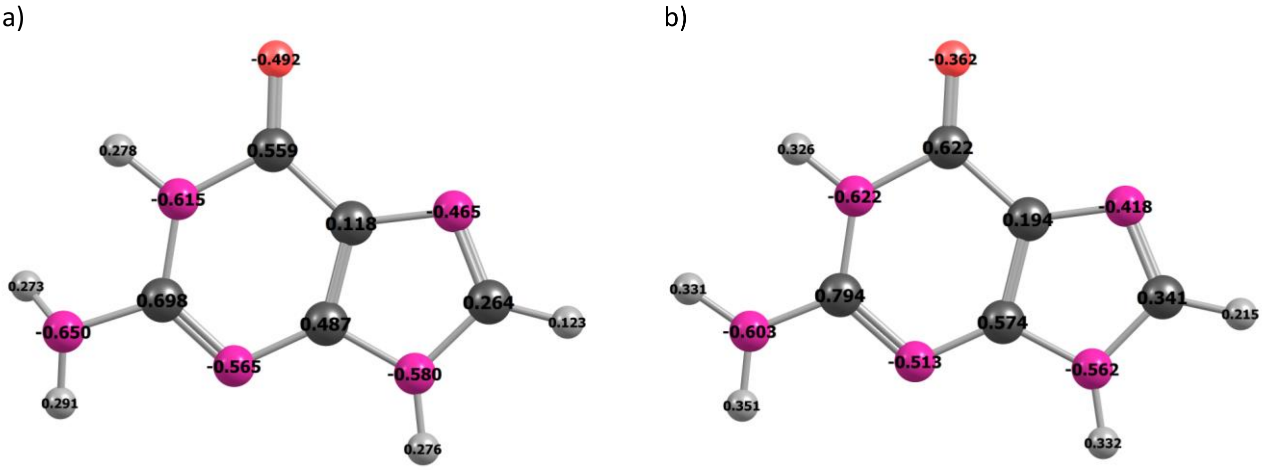


**Fig. S5** Mulliken charges in the neutral (a) and cation (b) states of the guanine nucleobase

### S4. Pathway from 4Fe4S to tRNA

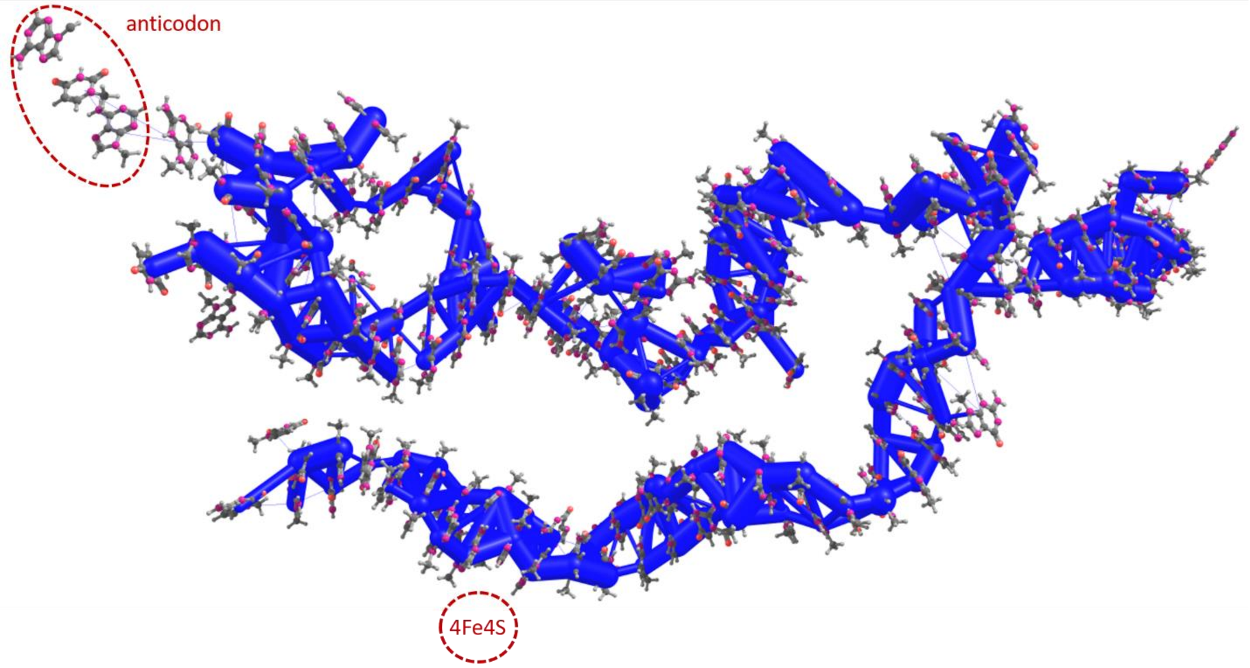


**Fig. S6** Hole transfer integrals *J* within the pathway from 4Fe4S to tRNA. The thickness of the cylinders represents the magnitude of *J* in the logarithmic scale; the largest *J* amount to 160 meV, the smallest drawn are of the order of 10 meV.
